# Joint bacterial traces in the gut and oral cavity of Colitis patients provide evidence for saliva as rich microbial biomarker source

**DOI:** 10.1101/2025.03.17.643500

**Authors:** Jacqueline Rehner, Madline Gund, Sören L. Becker, Matthias Hannig, Stefan Rupf, Jörn M. Schattenberg, the IMAGINE consortium, Verena Keller

## Abstract

The human microbiome, distributed across various anatomical sites, holds promise for identifying diagnostic biomarkers and therapeutic targets in disease. In inflammatory bowel disease (IBD), including ulcerative colitis (UC), interactions between the gut and oral microbiomes are crucial for understanding disease mechanisms and guiding interventions. The IMAGINE study sequenced 1,931 specimens from saliva, plaque, stool, and other sources in patients and healthy controls. Here, we assess whether the oral (saliva/plaque) or gut microbiota provides greater diagnostic potential in IBD and examine shared dysregulation across sample types. Among 177 oral samples (102 healthy, 75 IBD) and 92 stool samples (57 healthy, 35 IBD), we identified 240 distinct strains in plaque, 229 in saliva, and 231 in stool, with 46 strains present in all three. Saliva showed a significantly higher average effect size (0.2) than stool (0.04) and plaque (0.06). Notably, *Actinomyces* sp., *Bifidobacterium dentium*, and *Veillonella parvula* exhibited increased effect sizes, suggesting their potential as diagnostic markers or therapeutic targets. These findings indicate that microbiome profiling in IBD may improve diagnostics and treatment strategies.

## INTRODUCTION

The human microbiome has emerged as a cornerstone of modern medical research, offering profound insights into human health and disease. This intricate community of microorganisms, including bacteria, fungi, archaea, and viruses, resides across various body sites and influences numerous physiological processes, from immune system regulation to metabolism. As our understanding of microbiome functionality grows, so does our knowledge of its diagnostic and therapeutic potential. Diseases once attributed solely to genetic or environmental factors are now being reexamined through the lens of microbiome dysbiosis. Among these, inflammatory bowel disease (IBD), encompassing Crohn’s disease (CD) and ulcerative colitis (UC), ranks high among the most widely studied entities. Especially with respect to its complex interplay between host genetics, environmental triggers, and microbial imbalances, and close connection between the microbes and the affected organ, the gut, IBD has emerged as one role model of microbiota research. Globally, IBD affects over 7 millions of individuals^1^, with its prevalence rising sharply in newly industrialized regions, underscoring the urgent need for early and precise diagnostic tools that can be facilitated by the microbial community composition^2,3^. Studies that describe the effect of pro– and prebiotics underscore the importance of the gut microbiome for health and diseases^4-6^.

Traditionally, stool samples were the primary focus of microbiome research, largely due to their accessibility and the wealth of microbial information they provide. Fecal samples have revealed significant dysbiotic patterns in IBD patients, including a reduction in short-chain fatty acid-producing bacteria such as *Faecalibacterium prausnitzii* and an increase in pro-inflammatory taxa like adherent-invasive *Escherichia coli*^3,7^. However, while fecal microbiota studies have elucidated important aspects of IBD pathophysiology, they represent only a fraction of the broader microbial landscape. Other easily accessible sample types, such as saliva and interdental plaque have gained attention for their potential to offer complementary diagnostic insights. Evidence suggests a key role of oral microbiota on metabolic and other diseases^8^. Moreover, a shift from the oral cavity to the gut with pro-inflammatory potential is well known^9^. The oral microbiome relates also to systemic conditions like Parkinson’s disease, where saliva-based analyses have shown promising diagnostic accuracy, often surpassing stool in early disease detection^10^. It is also known that the diseases periodontitis and diabetes are linked and have also an influence on the gut microbiome^11,12^. Similarly, rapidly developing sequencing technologies allow discovering pivotal information about host–microorganism interactions and health outcomes in general^13^. This underlines the utility of microbiota as a diagnostic reservoir and for therapeutic approach. These findings challenge the traditional stool-centric approach and advocate for a multi-sample strategy in microbiome diagnostics.

Most microbiome studies to date describe correlations between microbial community changes and disease states, providing valuable insights but often lacking mechanistic depth. To move beyond correlation and toward functional understanding, it is critical to investigate the metabolic and biosynthetic capabilities of the microbiota. One key area is the study of biosynthetic gene clusters (BGCs), which encode the molecular machineries responsible for producing secondary metabolites, also known as natural products. These compounds play essential roles in microbial communication, competition, and host interactions^14^. The human microbiome represents a vast, largely untapped reservoir of natural products with therapeutic potential, ranging from antibiotics to immunomodulators. Harnessing this resource could not only advance microbiome-based diagnostics, but also drive the discovery of novel bioactive compounds for medical applications. Understanding and exploiting the functional capabilities encoded within BGCs will be central to this next phase of microbiome research.

Following these two recent trends, i.e. (1) understanding the role of multiple metagenomes from patients and (2) considering the potential of patient-derived microbiota as natural producers, we initiated the IMAGINE study (Identification of microbial antibiotics to protect the physiologic microbiota at body surfaces). The IMAGINE study exemplifies this paradigm shift by employing a cross-disease, cross-sample analysis to explore microbial variations across different body sites and health conditions^15^. This comprehensive approach revealed notable site-specific differences in microbial composition and functionality, reinforcing the diagnostic value of integrating multiple sample types. Specifically, in the context of IBD, the study highlighted distinct microbial patterns across stool, saliva, and interdental plaque, offering a more nuanced understanding of disease-associated dysbiosis. Such findings underscore the need to move beyond single-sample studies to capture the complex interplay between microbial communities at different anatomical sites. Despite these advancements, significant challenges remain in microbiome research. One of the most pressing issues is the lack of standardized protocols for sample collection, storage, and processing. Variations in DNA extraction methods, for example, can lead to discrepancies in microbial diversity and composition, particularly affecting hard-to-lyse Gram-positive bacteria^16^. These methodological inconsistencies complicate cross-study comparisons and hinder the reproducibility of findings. Recognizing this, the IMAGINE study placed a strong emphasis on standardization, employing rigorously validated protocols to ensure data quality and comparability^17^. This methodological rigor is essential for translating microbiome research into clinical practice, where diagnostic tools must be both accurate and reproducible.

In analyzing the highly standardized data from the IMAGINE study we identified distinct patterns in the microbial composition within the oral samples (interdental plaque and saliva) and stool samples of patients diagnosed with IBD. The objective of this study was to ascertain whether the oral (saliva/plaque) or gut microbiota offer greater diagnostic potential in IBD patients. Additionally, we sought to determine whether dysregulation of several species occurs in both sample types. The current study details common grounds and important differences in this context informing future diagnostic and therapeutic developments in IBD (***Fig. 1A***).

**Figure 1.**
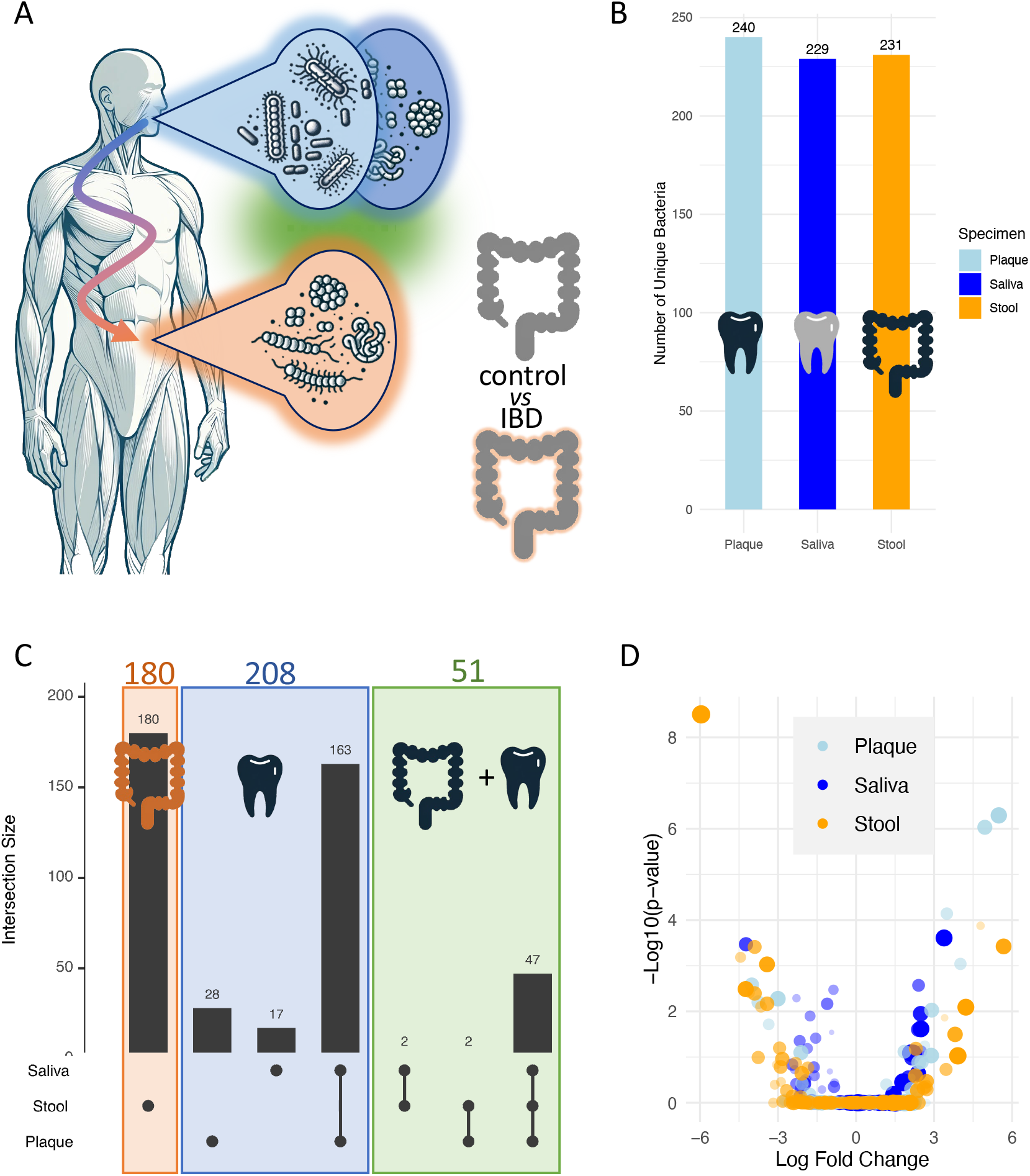
**(A)** Study set up. We set to compare gut and oral cavity microbiomes to find joint traces between the two body sites. **(B)** Number of species detected in the three sample types. (**C)** Upset-plot that provides an overview how the species discovered in the three sample types are distributed. **(D)** Volcano plot that presents the log fold changes versus the p-values for diagnosing Colitis. Each dot corresponds to one bacterial species in one of the three sample types. The sample type is represented by the different colors.

## METHODS

### Clinical sampling

Detailed procedures are described in the IMAGINE main manuscript^15^. In brief, clinical samples were collected from participants at Saarland University Medical Center in Homburg, Germany, following written informed consent. Ethical approval for the study was granted by the ethics committee of the local medical association (Ärztekammer des Saarlandes, ID: 131/20). Each participant underwent a thorough medical and dental examination to identify diseases of interest, and an in-depth medical history was recorded, including relevant factors such as medication, diet, activity levels, smoking, alcohol consumption, and co-morbidities. Samples included saliva, interdental plaque, conjunctiva swabs, throat swabs, stool, and skin swabs from the forehead and arm, as well as affected areas for participants with Acne inversa. The diagnostic procedures and sample collection were performed as described in the original study (Quelle 7). Here, we provide a summary limited to the sample types relevant to this manuscript. Stool samples (500 mg to 1 g) were collected using a paper toilet-hat and sterile collection tubes with integrated spoons. Interdental plaque was obtained by brushing three interdental spaces in each quadrant using 12 micro applicators, which were then transferred to an ESwab tube containing Amies Medium. Saliva samples were gathered in sterile falcon tubes by having participants spit unstimulated saliva for 5 minutes. All collected swabs and samples were immediately placed in the appropriate transport medium and frozen at –80 °C for preservation.

### DNA extraction, library prep and sequencing

Detailed DNA extraction and sequencing protocols are described in the IMAGINE main manuscript^15^. In brief, DNA extraction was performed using the Qiagen QiAamp Microbiome Kit (Qiagen, Hilden, Germany) following the manufacturer’s protocol. Swabs were vortexed in 1.5 ml Amies Medium for 2 minutes to release the microbial mass, which was then processed for DNA extraction. For fecal samples, 250 mg of stool was mixed with 500 µl of buffer AHL prior to DNA extraction. Interdental plaque collected via micro applicators was vortexed rigorously for 2 minutes in 1 ml PBS (pH 7.4), and the resulting bacterial suspension was used for extraction. Saliva samples were homogenized by brief vortexing, and 1 ml of saliva was processed directly. Mechanical cell disruption was achieved using the MP Biomedicals™ FastPrep-24™ 5G Instrument (FisherScientific GmbH, Schwerte, Germany) set at 6.5 m/s for 45 seconds, repeated twice with 5-minute intervals on ice between cycles. Extracted DNA was eluted into 50 µl of buffer and quantified using a NanoDrop 2000/2000c spectrophotometer (ThermoFisher Scientific, Wilmington, DE) for microvolume UV-Vis measurements^30^. Extracted DNA from all samples was sent to Novogene Company Limited (Cambridge, UK) for metagenomic library preparation and paired-end (PE150) sequencing using the Illumina HiSeq platform. To ensure high-quality data, sparsely collected biospecimens (n=47), substandard samples (n=1304), and anomalous samples (n=201) were excluded from the analysis.

### Primary data analysis

Detailed primary data analysis procedures are described in the IMAGINE main manuscript^15^. In brief, host read removal was done with KneadData^31^ and the human sra-human-scrubber^32^. Paired-end reads were only kept if none of the read pairs mapped to the human reference. After decontamination, we performed fastp^33^ QC and visualized results with MultiQC^34^. Mash^35^ was applied for comparing MinHash distances. Embeddings were generated in R relying on the UMAP package^36^. To remove outliers, a Grubb’s test on the mean of all pairwise MinHash distances was done in an iterative manner.

MetaPhlAn3^31^ was applied to profile quality-controlled samples and relative counts were rescaled to absolute counts. To estimate alpha diversity, Shannon diversity was used. Differential abundance was computed using ANCOM-BC^37^. All p-values were adjusted based on Benjamini-Hochberg adjustment if not mentioned explicitly. In visualizations of abundances, absolute counts were center log ratio normalized. SPAdes^38^ was used for metagenomic assemblies and the quality was assessed with QUAST ^39^. Scaffolds were binned with MetaBAT2^40^. MAGs were aggregated and dereplicated across samples with dRep^41^. Only mid and high-quality genomes were kept based on MIMAG’s^42^. To assess the novelty of the SGBs, GTDB r214^43^, the Unified Human Gastrointestinal Genome collection^44^, the Singapore Platinum Metagenomes Project^45^, and Pasolli et al.^46^ with Mash distances <= 0.05 were used, followed by validation with FastANI^47^.

### Downstream data analysis

Downstream bioinformatics analysis was conducted to extract and visualize meaningful insights from the microbial metagenomic data that we extracted from IMAGINE. The dataset consisted of taxonomic, statistical, and metadata information for various specimens, including stool, saliva, and interdental plaque. Altogether, 64,588 disease associations between different specimen types and diseases were processed. Of those, we considered 621 associations between colitis (the dominant IBD group in IMAGINE) and stool, saliva or plaque. Data preparation, visualization, and statistical evaluation were carried out using R, with detailed steps outlined below. Initially, IMAGINE data were imported and preprocessed. The dataset was read as a matrix, with separate annotation and numerical matrices created for metadata (e.g., specimen types, taxonomic species) and statistical values (e.g., p-values, fold changes, effect sizes). Taxonomic species were extracted from the hierarchical bacterial OTU column, isolating the species-level designation. Filtering focused on samples flagged as “all” in the comparison column, excluding other entries. Volcano plots were generated to visualize the relationship between log-fold changes and p-values for specific group comparisons, such as Colitis versus Healthy. Specimen types of interest – stool, saliva, and plaque – were highlighted using distinct color schemes. Scatterplots compared effect sizes across specimen types at the species level with regression lines quantifying the relationships between effect sizes in saliva versus plaque, saliva versus stool, and plaque versus stool. For species present across all three specimen types, dot plots were created. These displayed effect sizes for each specimen type, with color coding and point sizes indicating effect size magnitude. Species were ordered by their average effect size across specimen types, highlighting dominant taxa. Back-to-back histograms are generated to illustrate the distribution of effect sizes for plaque, saliva, and stool. Similarly, heatmaps visualized –log10-transformed p-values for significant bacterial species, categorized by specimen type. For computing beeswarm plots with significance values the ggbetweenstats package has been used^48^. Lastly, UpSet plots depicted the overlap of species presence across stool, saliva, and plaque, providing an overview of shared and unique taxonomic features.

### PubMed/MEDLINE Search

To assess the plausibility of our findings, we conducted a comprehensive PubMed/MEDLINE search. Each of the 38 species identified across all three sample types was queried using a combination of terms: “oral,” “plaque,” “saliva,” “stool,” and “gut” (representing five specimen types), paired with “colitis” and “IBD” (representing two diseases). This approach yielded a total of 380 individual queries, systematically designed to evaluate the associations between species, specimen types, and disease contexts through an evidence-based review of existing literature.

## RESULTS

We identified 700 species-to-disease associations across the three specimen types. These associations were distributed evenly among the specimen types: interdental plaque samples showed the highest number with 240 associations, followed by stool with 231, and saliva with 229 (***Fig. 1B***). Importantly, these 700 associations do not correspond to 700 distinct species, as some species were associated with diseases in more than one specimen type. In total, the 700 associations represented 439 unique species. Of these, 225 species were specific to a single specimen type, 167 were shared between any two specimen types, and 47 species (10.7%) were identified in all three habitats. When stratified by anatomical region, 208 species were detected exclusively in the oral cavity, while 180 were specific to stool. An additional 57 species were shared between the oral cavity and stool (***Fig. 1C***). This overlap highlights the interplay between different microbiome niches and their role in disease associations. Given the focus on IBD, particularly UC as the predominant subtype in IMAGINE, we focused our analysis on 621 associations between the three specimen types and UC. Volcano plots comparing fold changes and p-values revealed intriguing patterns: significant negative associations were predominantly observed in stool samples, while significant positive associations were more frequent in saliva samples (***Fig. 1D***). A detailed analysis revealed several notable bacterial associations across different specimen types. In stool samples, *Gemmiger formicilis* emerged as particularly significant, with a strong negative effect size of –0.81 (p = 3.13 × 10^−9^), suggesting a marked reduction in its abundance under the studied conditions. *Clostridium innocuum* also showed a smaller positive effect size of 0.06 (p = 1.33 × 10^−4^), indicating subtle yet significant changes in its abundance. In plaque samples, *Bifidobacterium dentium* exhibited a positive effect size of 0.46 (p = 9.28 × 10^−7^), underscoring the unique microbial composition associated with this specimen type, while *Olsenella profusa* displayed a moderate positive effect size of 0.26 (p = 7.22 × 10^−5^), supporting its relevance within the microbial community of this niche. In saliva samples, *Propionibacterium acidifaciens* demonstrated consistent associations, appearing in both plaque and saliva with positive effect sizes of 0.59 (p = 5.10 × 10^−7^) and 0.66 (p = 2.46 × 10^−4^), respectively, highlighting its potential role in these environments. Conversely, *Lachnospiraceae bacterium oral taxon 096* exhibited a notable negative effect size of –0.40 (p = 3.39 × 10^−4^), suggesting a decrease in its presence. While these findings underscore the diverse and dynamic microbial changes occurring in stool, plaque, and saliva, they highlight the importance of including multiple specimen types to obtain a comprehensive picture of microbiome-disease interactions. We thus initiated a direct comparison between the gut and oral cavity.

Because p-values can be misleading^18^ (e.g. they do not convey the magnitude of an effect, can be influenced by sample size, and are susceptible to misinterpretation) and fold changes can likewise be misleading (e.g. due to their sensitivity to variability in data and potential to overemphasize minor differences) we have chosen to focus on effect sizes, which provide a more balanced and interpretable measure of the strength of associations, offering a clearer understanding of the practical significance of our findings. First, we computed back-to-back histograms to compare the effect sizes for diagnosing colitis samples in stool versus saliva (***Fig. 2A***) and stool versus interdental plaque (***Fig. 2B***). These histograms revealed a clear shift towards higher effect sizes in saliva compared to both, stool and interdental plaque. To further explore these differences, we utilized beeswarm plots to visualize the absolute effect sizes across the three specimen types. Statistical comparisons were conducted using the Games-Howell post hoc test, following Welch’s ANOVA. This test is particularly appropriate for scenarios with unequal group variances or differing sample sizes, ensuring robust pairwise comparisons. The analysis revealed that saliva exhibited notably higher effect sizes (mean of 0.2) overall compared to stool (mean of 0.04) and interdental plaque (mean of 0.06) (***Fig. 2C***). These findings underscore the distinctiveness of saliva in capturing biological effects relevant to UC and emphasize its potential as a diagnostic specimen. In the previous analyses, the direct relationships of individual bacterial species across specimen types were not explicitly addressed. To explore these relationships in greater detail, we computed scatter plots to compare the effect sizes of each species between pairs of specimen types. Specifically, we compared species effect sizes between saliva and interdental plaque (***Fig. 2D***), saliva and stool (**Fig. 2E**), and interdental plaque and stool (***Fig. 2F***). These scatter plots allow for a direct species-by-species assessment of the effect sizes, highlighting potential correlations and trends between the specimen types. The results demonstrate a strong positive correlation between saliva and interdental plaque, indicating a shared microbial signature in these oral cavity specimens. This suggests that species showing higher effect sizes in saliva are likely to exhibit similar trends in interdental plaque, reflecting their common origin within the oral microbiome. In contrast, when comparing saliva to stool, we observed no apparent correlation. This lack of association highlights the distinctiveness of the gut microbial profile compared to the oral cavity. Similarly, for interdental plaque versus stool, there is at most a slight positive correlation, which may reflect the influence of shared bacterial taxa between these two sites but with significant variability. These findings underline the strong divergence of microbial patterns between the oral cavity and the gut, as evidenced by the weak correlations in effect sizes between stool and oral samples. This divergence reinforces the importance of considering specimen-specific factors when analyzing microbial data and selecting specimen types for diagnostic purposes. Finally, we aimed to directly compare bacterial species that occur across all three specimen types. This approach provides a focused analysis of species with broad representation across different microbiomes, allowing for a more integrated perspective. In total, we identified 39 of such species. Among these, one entry corresponds to an unidentified species, included for the sake of completeness. Excluding this single exception, the analysis concentrated on 38 well-characterized bacterial species. This targeted approach provides valuable insights into the dynamics of these species and their potential relevance for understanding microbial patterns in relation to disease.

**Figure 2.**
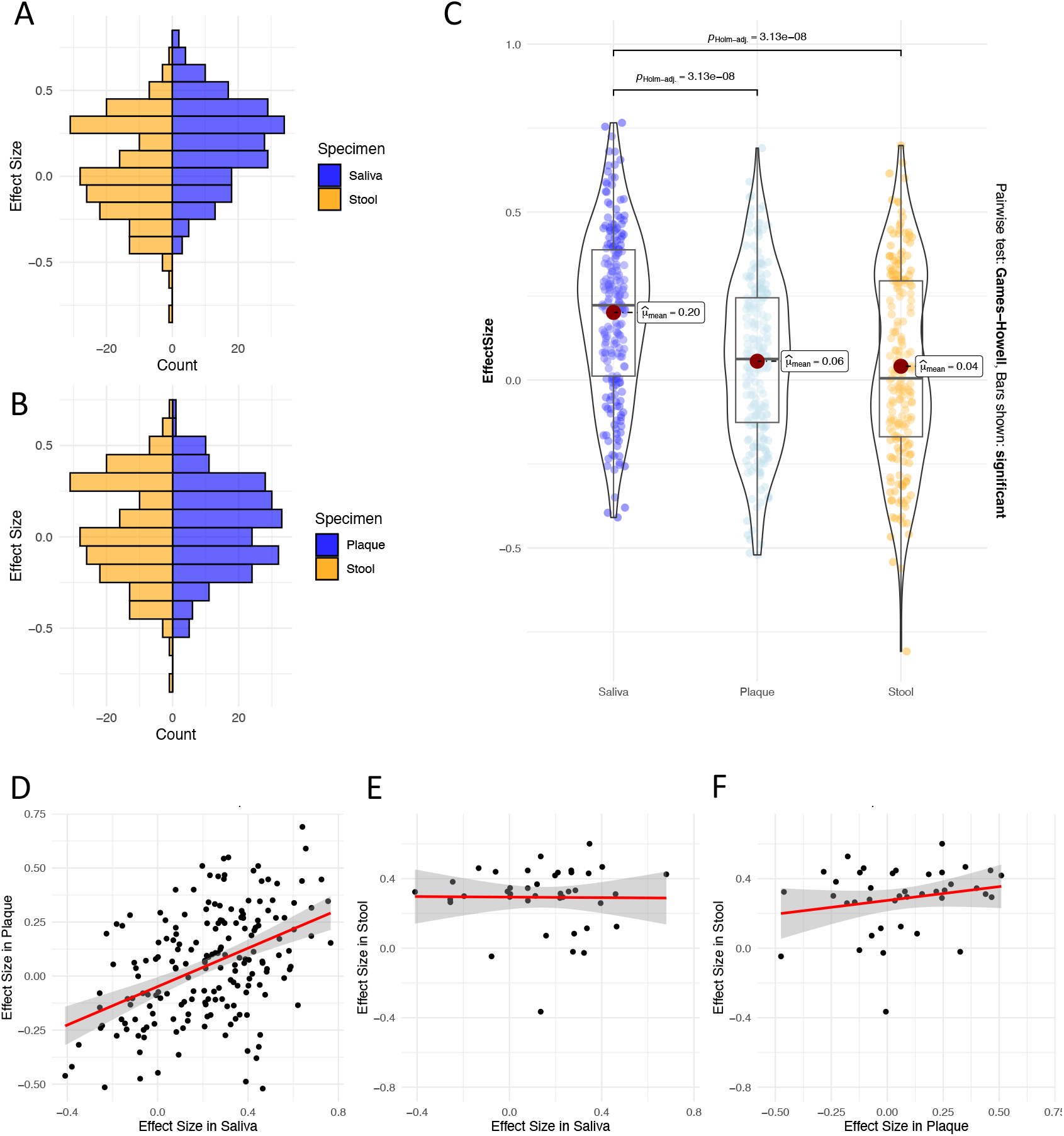
**(A)** Back-to-back histogram comparing the effect sizes between saliva (blue) and stool (orange) species. **(B)** Back-to-back histogram comparing the effect sizes between plaque (blue) and stool (orange) species. **(C)** Direct comparison of the effect sizes as beeswarm plots. Mean values and significance values are used to annotate the beeswarm plot and box plots per distribution are shown in the background for each sample type. **(D)** Scatter plot comparing the effect sizes in saliva to those in plaque. The regression is presented as red lines. The grey area represents the respective confidence interval. **(E)** Same panel as in (D) but for saliva and stool. **(F)** Same panel as in (D) but for plaque and stool.

The analysis of species present in all three specimen types reveals interesting patterns in effect sizes (**Fig. 3**) that warrant further consideration. Among the species identified, notable differences emerge based on their specimen-specific and overall effect sizes. The analysis revealed that *Veillonella parvula* exhibits the highest average effect size (0.43) across specimen types, driven by a markedly high effect size in saliva (0.68). This is accompanied by moderate contributions from stool (0.43) and interdental plaque (0.19). In contrast, *Gemella sanguinis*, with an average effect size of 0.41, demonstrates a more balanced distribution of effect sizes, with stool (0.47) being its highest, followed by saliva (0.40) and interdental plaque (0.35). Other species, such as *Actinomyces sp. oral taxon 448*, also stand out with a high average effect size (0.40). While interdental plaque (0.44) and saliva (0.46) contribute significantly to this value, the stool effect size is relatively lower (0.31). Conversely, *Bifidobacterium dentium* maintains a balanced distribution of high effect sizes across stool (0.45), interdental plaque (0.46), and saliva (0.27), leading to an average of 0.39. Interestingly, *Streptococcus parasanguinis* also shows a notable average effect size (0.38), with stool (0.42) and interdental plaque (0.51) contributing prominently, compared to a lower effect size in saliva (0.20). This indicates a potential specimen-specific relevance of this species for certain conditions. Overall, the stool samples consistently exhibit higher effect sizes for many species compared to saliva and interdental plaque. This is particularly evident for *Actinomyces sp. HPA0247* and *Actinomyces sp. oral taxon 181*, both of which have their highest individual effect sizes in stool. These findings contrast with the general trend of saliva showing the highest overall effect sizes, suggesting specific ecological or pathological roles for certain species in stool samples. These results not only highlight key species which are consistently associated with significant effect sizes but also emphasize the importance of specimen specific differences in microbial associations. These findings could guide targeted investigations into the role of these species in disease and health.

**Figure 3.**
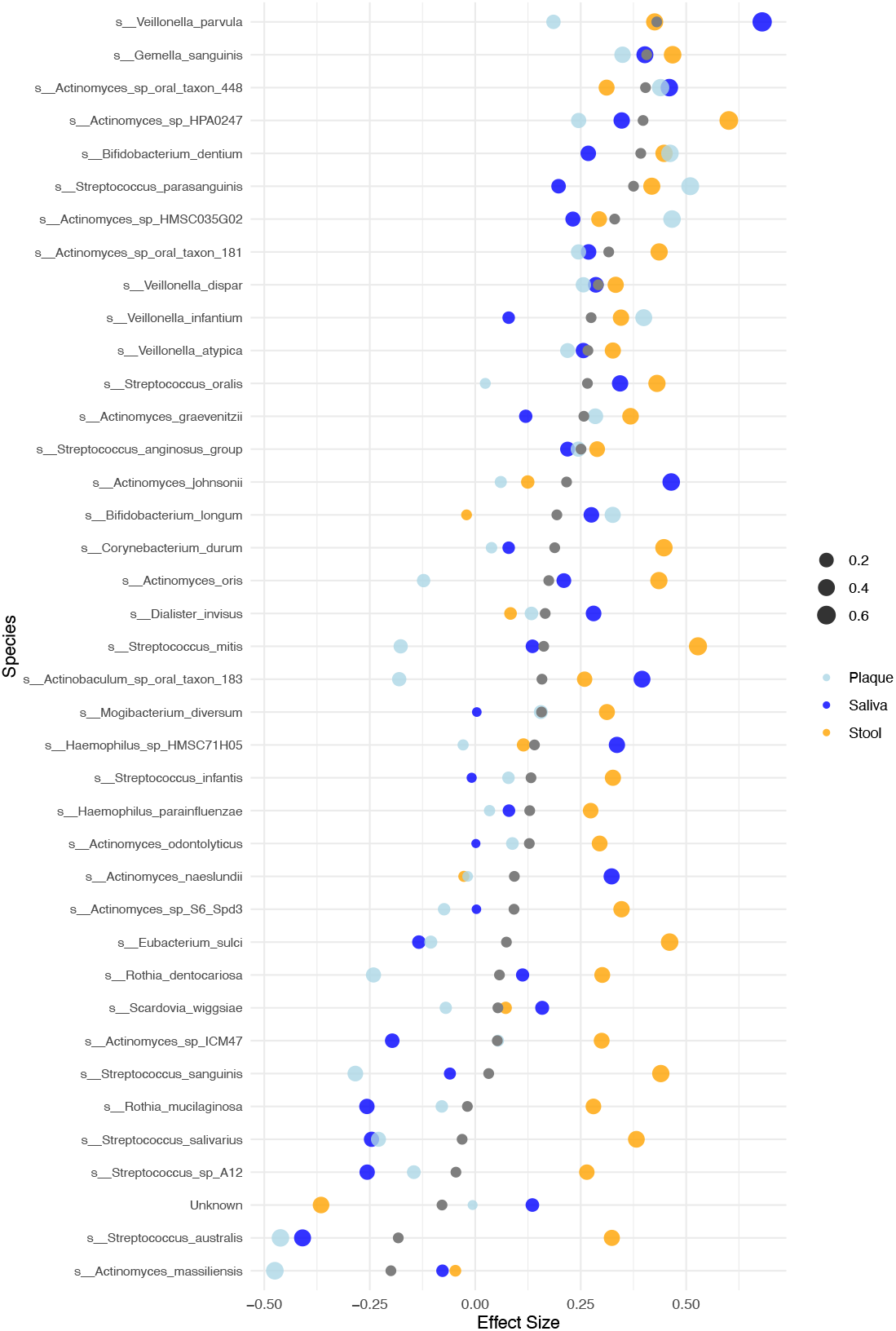
Effect sizes of bacterial species across saliva, plaque, and stool specimens. Each dot represents the effect size of a bacterial species in one of the three specimen types (saliva: blue, plaque: light blue, stool: orange). Gray dots indicate the average effect size of each species across all three specimen types. The species are ordered by their average effect size in ascending order. The size of the colored dots reflects the magnitude of the effect size. This visualization highlights the variability and trends in bacterial species’ contributions across the different specimen types.

The systematic PubMed/MEDLINE search described in the Methods section provided a comprehensive framework to evaluate potential associations between microbial species, specimen types, and disease contexts. Of the 380 queries performed, 50 returned at least one PubMed hit, offering evidence for the respective species, specimen type, and disease combination. In 13 cases, at least three supporting manuscripts were identified (Fig. 4A). *Bifidobacterium longum* emerged as the species with the strongest evidence, showing prominent associations with “gut” and “colitis” (56 hits). This species also demonstrated significant associations with “stool” and “colitis” (26 hits), “gut” and “IBD” (25 hits), as well as “oral” and “colitis” (20 hits). These findings highlight its consistent presence across multiple specimen types and its relevance to gastrointestinal and systemic conditions. *Streptococcus salivarius* also showed notable associations, particularly with “oral” and “colitis” (4 hits), “stool” and “colitis” (4 hits), and “gut” and “colitis” (3 hits), underscoring its role as a commensal in the oral microbiome. Other species, such as *Haemophilus parainfluenzae*, were linked to “gut” and “colitis” (3 hits), while *Dialister invisus* and *Veillonella dispar* demonstrated associations with “gut” and “IBD” (3 hits each). Additionally, *Bifidobacterium dentium* was identified in the context of “gut” and “colitis” (3 hits), illustrating the diversity of microbial species implicated in these diseases.

**Figure 4.**
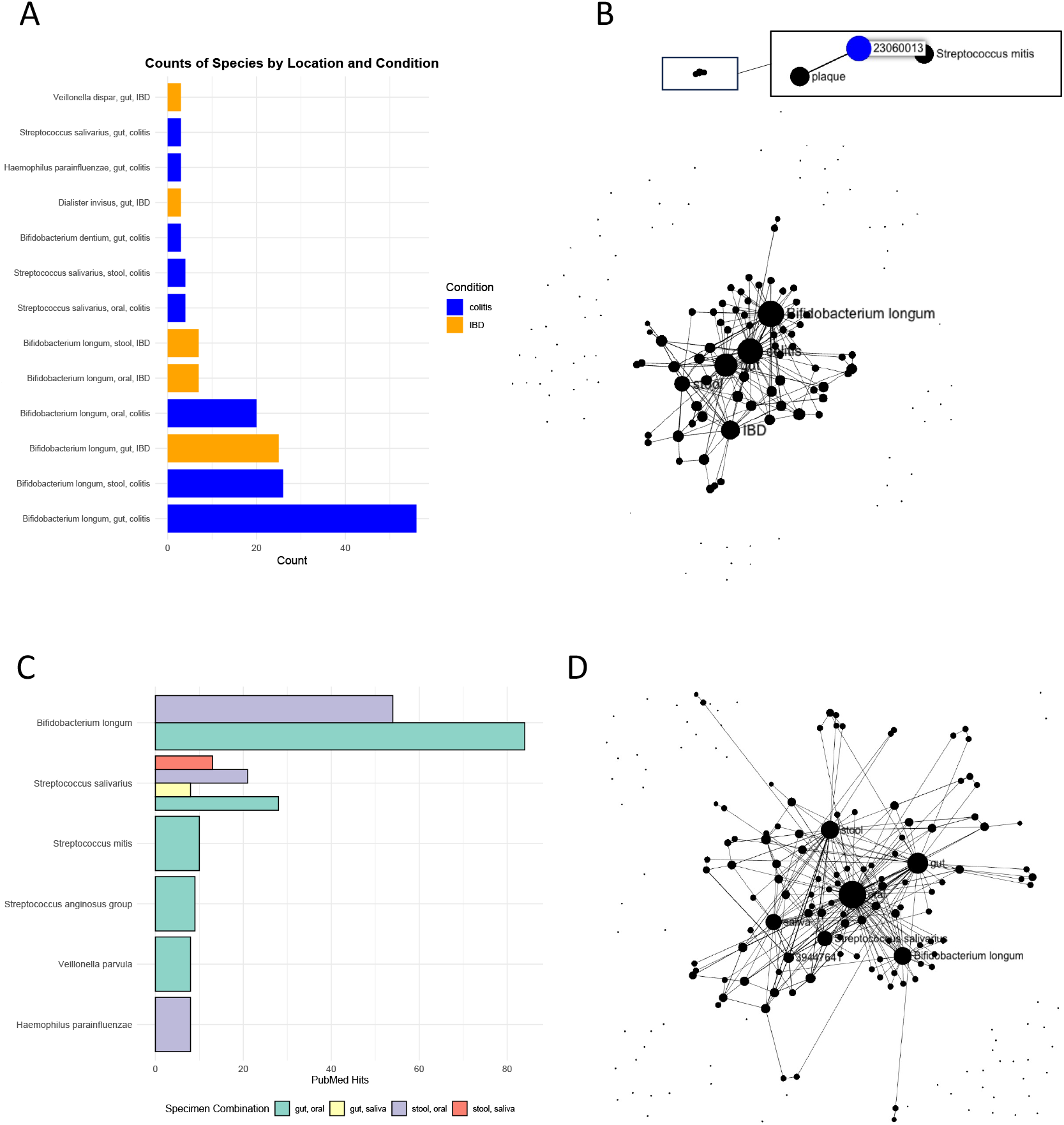
**(A)** Hits for the PubMed search of the 38 species with respect to the specimen types and colitis and IBD. Colors represent the disease and the bar length the number of PubMed hits. Only combinations with at least three hits are presented. **(B)** Publication network. The network connects search results to manuscripts. It shows the most relevant hubs represented by larger nodes. In this case, *Bifidobacterium longum*, IBD, colitis and stool were the most frequent hits. **(C)** Similar to the results in panel (A) we carried out a literature search for the 38 species containing both, gut or stool and saliva or oral, independent on diseases. Besides *Bifidobacterium longum, Streptococcus salivarius* shows up as relevant diagnostic species. **(D)** Publication network as presented in panel (B) for the disease agnostic search.

To gain a more comprehensive understanding, we visualized the results as a network where the different queries are linked to their respective manuscripts, represented as dots connected by lines (***Fig. 4B***). This network provides an intuitive overview of the associations between bacterial species, specimen types, and the literature evidence supporting them. The network confirmed *Bifidobacterium longum* as the dominant species, with its associations spanning multiple specimen types and a large number of manuscripts. Similarly, *Streptococcus salivarius* emerged as a hub node, indicating its significant relevance across different environments and studies. In addition to these well-connected hub nodes, one distinct combination stood out: *Streptococcus mitis* in plaque^19^. This manuscript provides evidence that, despite similar clinical periodontal parameters, IBD patients exhibit elevated levels of bacteria associated with opportunistic infections in inflamed subgingival sites. These findings prompted us to broaden our analysis beyond the context of colitis or IBD specifically, to explore whether similar bacterial profiles and associations occur across different conditions and environments. We expanded our analysis by performing a PubMed search to evaluate whether the identified bacterial species are commonly associated with other diseases, irrespective of their link to UC or IBD. Specifically, we queried each species for the combinations of “gut,” “stool,” “oral,” and “saliva” in a disease-agnostic manner. From these searches, we identified significant associations between species and specific specimen types, with 10 species/specimen combinations hit counts (***Fig. 4C***). Among these, *Bifidobacterium longum* demonstrated the highest number of hits, with 84 articles linking it to the “gut, oral” combination, followed by 54 hits for its association with “stool, oral.” This finding emphasizes the broad distribution and potential clinical importance of *Bifidobacterium longum* across these environments. Importantly, as *Bifidobacterium longum* is widely recognized as a beneficial probiotic and commonly found in healthy individuals, its presence should not be misinterpreted as pathogenic. Instead, its frequent detection in both gut and oral samples, including in healthy controls, underscores its role in maintaining microbial balance rather than indicating disease association. *Streptococcus salivarius* was another prominent species, identified in multiple combinations, including 28 hits for “gut, oral,” 21 hits for “stool, oral,” and 13 hits for “stool, saliva.” Its widespread occurrence highlights its relevance in both gut and oral environments and suggests a versatile ecological niche. Other noteworthy findings include *Streptococcus mitis* with 10 hits for “gut, oral” and *Streptococcus anginosus* group with 9 hits for the same combination, underscoring the importance of these species in the gut-oral axis. Additionally, *Haemophilus parainfluenzae* and *Veillonella parvula* each garnered 8 hits for “stool, oral” and “gut, oral,” respectively, further reflecting the cross-compartmental presence of these bacteria (***Fig. 4D***). These results suggest the biological relevance of our findings, as many of the identified species have previously been reported in association with the studied specimen types and diseases. This alignment with existing literature supports the consistency of our analysis while also highlighting key microbial species that warrant further investigation.

## DISCUSSION

Human microbiota is increasingly recognized as a key player in maintaining health and contributing to disease^20^. Dysbiosis, or microbial imbalance, has been implicated in various conditions, including IBD and its subtypes such as UC^2^. While the gut microbiota has been extensively studied in this context, other body sites, such as the oral cavity, are gaining attention for their potential roles in systemic and localized inflammation^21^. This study aimed to comprehensively evaluate bacterial contributions to UC across multiple specimen types, focusing on saliva, interdental plaque, and stool, to capture the diversity and specificity of microbial shifts. Our analysis revealed notable differences in microbial effect sizes across specimen types. Saliva consistently exhibited higher effect sizes, suggesting it might serve as a sensitive and non-invasive proxy for detecting microbial alterations associated with UC. Stool samples, while traditionally used for gut microbiome studies, showed unique species with high effect sizes, indicating a distinct microbial signature reflective of gut-specific dysbiosis. Plaque, on the other hand, presented generally lower effect sizes, but with notable exceptions that emphasize the relevance of the oral microbiota in disease mechanisms. Importantly, factors such as dental health status, including the presence of caries or periodontal disease, may also influence microbiome composition and should be considered when interpreting microbial shifts across specimen types.

We find it evident that highlighting the potential of saliva as a source for diagnostic microbial signatures is important. But we hypothesize that the patterns of interaction between oral and gut microbiota extend beyond IBD and indicate a broader role in systemic health and disease, as highlighted by multiple recent studies. Notably, the convergence of evidence from diverse conditions, including liver function recovery, cardiovascular diseases, and hepatic encephalopathy, emphasizes the intricate relationship between the microbiota of different body sites and systemic health^22,23^. *Bifidobacterium longum*, a key gut microbe, has been implicated in postoperative liver function recovery for hepatocellular carcinoma (HCC). A clinical trial demonstrated that oral administration of a probiotic cocktail containing *B. longum* reduced delayed recovery rates, shortened hospital stays, and improved overall survival. These benefits were linked to diminished liver inflammation, reduced fibrosis, and hepatocyte proliferation, underscoring the therapeutic potential of gut microbiota modulation beyond IBD^24^. Similarly, *Haemophilus parainfluenzae*, typically associated with the oral cavity, has been identified as a modifier of Crohn’s disease (CD) progression. Its presence in the gut, likely facilitated by periodontal disease, correlates with increased intestinal inflammation mediated through strain-specific interactions with host immune pathways. This highlights a pathogenic mechanism wherein oral bacteria exacerbate intestinal inflammation, suggesting a potential route for oral-to-gut microbial transmission^25^. The connection between oral and gut microbiota is further demonstrated in cardiovascular diseases such as acute myocardial infarction (AMI). Oral species, including *Streptococcus oralis, Streptococcus parasanguinis*, and *Streptococcus salivarius*, have been shown to colonize the gut and aggravate AMI in murine models. These findings suggest that monitoring and controlling specific oral bacteria could represent a novel therapeutic avenue for cardiovascular diseases^26^. On the other hand, some findings highlight the complexity and potential controversies in interpreting the role of microbiota. For instance, in untreated chronic periodontitis among IBD patients, CD patients were found to harbor higher levels of *Streptococcus anginosus, Streptococcus mutans*, and other opportunistic pathogens compared to both UC patients and healthy controls. Notably, these bacteria are commonly associated with dental caries, suggesting either a direct link between CD and caries or an underlying dietary pattern—potentially characterized by higher carbohydrate intake—that distinguishes CD patients from both UC patients and healthy individuals. These findings indicate that despite a similar clinical presentation of IBD, the microbiota composition varies significantly between subgroups, potentially influencing disease progression and associated comorbidities^19^.

In interpreting our data, it is essential to consider the complexity arising from the dynamic nature of the microbiome, which can be influenced by factors such as diet^27^, medication^28^, and disease state. For instance, exclusive enteral nutrition (EEN), a first-line therapy for pediatric CD, has been shown to induce significant changes in gut microbiota, including shifts in strain-level dynamics and metabolite profiles^29^. Such findings emphasize the microbiome’s responsiveness to therapeutic interventions and its potential role in personalized medicine. However, they also underscore the need for longitudinal studies to account for temporal variations in microbial composition and function. In addition to its diagnostic potential, the microbiome offers valuable insights into disease mechanisms. Multi-omics approaches integrating metagenomics, metabolomics, and other datasets have shed light on the intricate relationships between microbial taxa and host metabolic pathways. For example, a cross-cohort analysis revealed consistent alterations in gut microbial species and metabolite profiles associated with IBD, providing a robust framework for biomarker discovery^2^. Such integrative analyses are particularly relevant for complex diseases like IBD, where multiple factors converge to drive pathogenesis. Beyond those challenges, it is important to acknowledge limitations of our study. One key limitation is the cohort, which may not fully represent the variability of microbiota compositions across diverse populations or account for significant interindividual differences. Larger and more diverse datasets, encompassing tens of thousands of samples, are crucial to validate our findings and explore broader implications. Moreover, our analysis was complicated by uneven group sizes across different specimen types and patient groups. To address this limitation and ensure robust biological interpretation, we prioritized effect sizes over strict statistical significance thresholds (e.g., p-values). Additionally, while our cross-disease approach allowed us to identify common microbial patterns, it may have oversimplified distinct mechanisms underlying specific diseases. The role of comorbidities, which can significantly influence microbiota profiles and disease trajectories is another grant challenge in respective studies. Moreover, future studies with a longitudinal design are essential to uncover how microbiota changes dynamically over time and how these changes correlate with disease progression or remission. To overcome limitations, we put our results in the context of the literature. But the reliance on PubMed searches for validation introduces its own set of challenges. While useful for contextualizing findings within existing literature, search algorithms, inaccurate and misleading string matching, incomplete indexing, and publication biases may have influenced the results. We potentially excluded relevant studies or overemphasized certain patterns. This limitation underscores the importance of complementing literature-based approaches with experimental validation to ensure robustness. Additionally, our analysis focused on identifying key microbial species, but the potential role of low-abundance species and complex microbiota-host interactions might have been overlooked due to the resolution of the analytical methods used. Finally, the observational nature of this study precludes definitive conclusions about causality. Experimental validation, such as animal models or clinical interventions, will be critical to confirm the biological significance of the identified microbial associations. Taken together, these limitations highlight the importance of cautious interpretation and pave the way for future research to address these gaps systematically.

Building on known insights and considering known limitations, this study aims to systematically analyze and compare the metagenomes of stool, saliva, and plaque samples from IBD patients. By leveraging high-throughput sequencing and rigorous multi-sample integration, we seek to identify disease-specific microbial signatures and evaluate their diagnostic accuracy. This work not only addresses the limitations of single-sample studies but also contributes to the growing body of evidence supporting the use of multi-sample approaches in microbiome diagnostics. Ultimately, we aspire to advance the field toward a more comprehensive and personalized understanding of IBD, paving the way for improved diagnostic and therapeutic strategies.

## Acknowledgements

Our appreciation extends to all individuals associated with Saarland University especially the Center for Bioinformatics, Saarland University Medical Center, and Helmholtz Institute for Pharmaceutical Research Saarland, whose contributions significantly enriched this study. Furthermore, we extend our gratitude to the participants of this study, whose consent allowed us to acquire and scrutinize clinical samples, thereby unravelling insights into their intricate microbial composition and functions.

## Data Availability

The raw metagenomic sequencing data, after the removal of ambient human DNA, has been deposited in the Sequencing Read Archive (SRA) under the accession code PRJNA1057503: https://www.ncbi.nlm.nih.gov/bioproject/PRJNA1057503

